# Empirical assessment of published effect sizes and power in the recent cognitive neuroscience and psychology literature

**DOI:** 10.1101/071530

**Authors:** Denes Szucs, John PA Ioannidis

## Abstract

We have empirically assessed the distribution of published effect sizes and estimated power by extracting more than 100,000 statistical records from about 10,000 cognitive neuroscience and psychology papers published during the past 5 years. The reported median effect size was d=0.93 (inter-quartile range: 0.64-1.46) for nominally statistically significant results and d=0.24 (0.11-0.42) for non-significant results. Median power to detect small, medium and large effects was 0.12, 0.44 and 0.73, reflecting no improvement through the past half-century. Power was lowest for cognitive neuroscience journals. 14% of papers reported some statistically significant results, although the respective F statistic and degrees of freedom proved that these were non-significant; p value errors positively correlated with journal impact factors. False report probability is likely to exceed 50% for the whole literature. In light of our findings the recently reported low replication success in psychology is realistic and worse performance may be expected for cognitive neuroscience.

## Introduction

Low power and selection biases, questionable research practices and errors favouring the publication of statistically significant results have been proposed as major contributing factors in the reproducibility crisis that is heavily debated in many scientific fields (Nosek et al. 2015a,b; Ioannidis et al. 2014a,b; Ioannidis 2005). Here, we aimed to get an impression about the latest publication practices in the closely related cognitive neuroscience and (mostly experimental) psychology literature. To this end we extracted more than 100,000 records of degrees of freedoms, t values, F values and p values from papers published between Jan 2011 to Aug 2014 in 18 journals. Journal impact factors ranged from 2.367 (Acta Psychologica) to 17.15 (Nature Neuroscience). The data allowed us to assess the distribution of published effect sizes; to estimate the power of studies; to estimate the lower limit of false report probability and to confirm recent reports about p-value reporting errors. The text-mining approach we used enabled us to conduct a larger power survey than classical studies.

Low power is usually only associated with failing to detect existing (true) effects and therefore with wasting research funding on studies which a priori have low chance to achieve their objective. However, low power also has two other serious negative consequences: it results in the exaggeration of measured effect sizes and it also boosts false report probability, the probability that statistically significant findings are false (Ioannidis 2005; Button et al. 2013; Pollard and Richardson, 1987; Berger and Sellke, 1987).

First, if we use Null Hypothesis Significance Testing (NHST) then published effect sizes are likely to be, on average, substantially exaggerated when most published studies in a given scientific field with low power (Button et al. 2013; Sedlmeier and Gigerenzer, 1989; Rossi 1990; Cohen 1962; Hallahan and Rosenthal, 1996; Note that NHST is amalgamation of Fisher's significance testing method and the Nyman-Pearson theory. However, the concept of power is only interpreted in the Nyman-Pearson framework. For extended discussion see Gigerenzer and Marewski, 2015; Gigerenzer et al. 1990) because even if we assume that there is a fix true effect size, actual effect sizes measured in studies will have some variability due to sampling error and measurement noise. Underpowered studies will be able to classify as statistically significant only the occasional large deviations from real effect sizes; conversely, most measured effects will remain under the statistical significance threshold even if they reflect true relationships (Schmidt, 1992; Sterling et al. 1995; Ioannidis, 2008; see **Supplementary Figure 1A**). Because many “negative” findings are not published, even metaanalyses may exaggerate effects. Effect size inflation is greater, when studies are even more underpowered.

Second, regarding the NHST framework the long run False Report Probability (FRP) can be defined the probability that the null hypothesis is true when we get a statistically significant finding. The long run True Report Probability (TRP) can be defined as the probability that the alternative hypothesis is true when we get a statistically significant finding. Computationally, FRP is the number of statistically significant false positive findings divided by the total number of statistically significant findings. TRP is the number of statistically significant truly positive findings divided by the total number of statistically significant findings. FRP and TRP can be computed by applying Bayes theorem (see **Supplementary Material Section 4** for details and note that the concepts of FRP and TRP do not exist in the NHST framework.). In a simple case FRP can be computed as

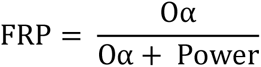

where O stands for H_0_:H_1_ odds and α denotes the statistical significance level which is nearly always α=0.05. So, for given values of O and α FRP is higher if power is low. As in practice O is very difficult to ascertain, high power provides the most straightforward ‘protection’ against excessive FRP (Ioannidis 2005; Button et al. 2013; Pollard and Richardson, 1987).

Because published effect sizes are likely to be inflated, it is most informative to determine the power of studies to detect pre-defined effect sizes. Hence, first we computed power from the observed degrees of freedom using supporting information from manually extracted records to detect effect sizes traditionally considered small (d=0.2), medium (d=0.5) and large (d=0.8) (Sedlmeier and Gigerenzer, 1989; Rossi 1990; Cohen 1962). Second, we also computed power to detect the reported effect sizes which were published across studies. Given that many of these effect sizes are likely to be inflated compared to the true ones (as explained above), this enabled us to estimate the lower limit of FRP (Ioannidis, 2005; Berger and Sellke, 1987). Further, we also examined p value reporting errors in the cognitive neuroscience literature as recent studies found that 12.5% to 20% of psychology papers have major p value reporting errors which affect judgments about statistical significance (Bakker and Wicherts, 2011; Nuijten et al. 2015; Veldkamp et al. 2014).

## Results and Discussion

The highly satisfactory performance of the extraction algorithm is reflected by the similarity between the distributions of automatically and manually extracted degrees of freedoms (which reflect the sample sizes of the studies, e.g. for an independent sample t-test, the degrees of freedom are the sample size minus 2) (**Fig. 1A**). During the validation process we assessed the proportion of one-sample, matched and two-sample t-tests which enabled us to use a mixture model to compute published effect sizes and power. The distribution of the computed effect sizes showed a near perfect match to the effect size distribution determined from records where D values were available which suggests that the mixture model we used is likely to well approximate the proportion of one-sample and two-sample t-tests (**Fig. 1B**). The computed D value distribution was slightly shifted to the right relative to the reported D value distribution, but both the medians (computed = 0.653; reported = 0.660) and means (computed = 0.938; reported=0.889) were very similar.

**Figure 1.**
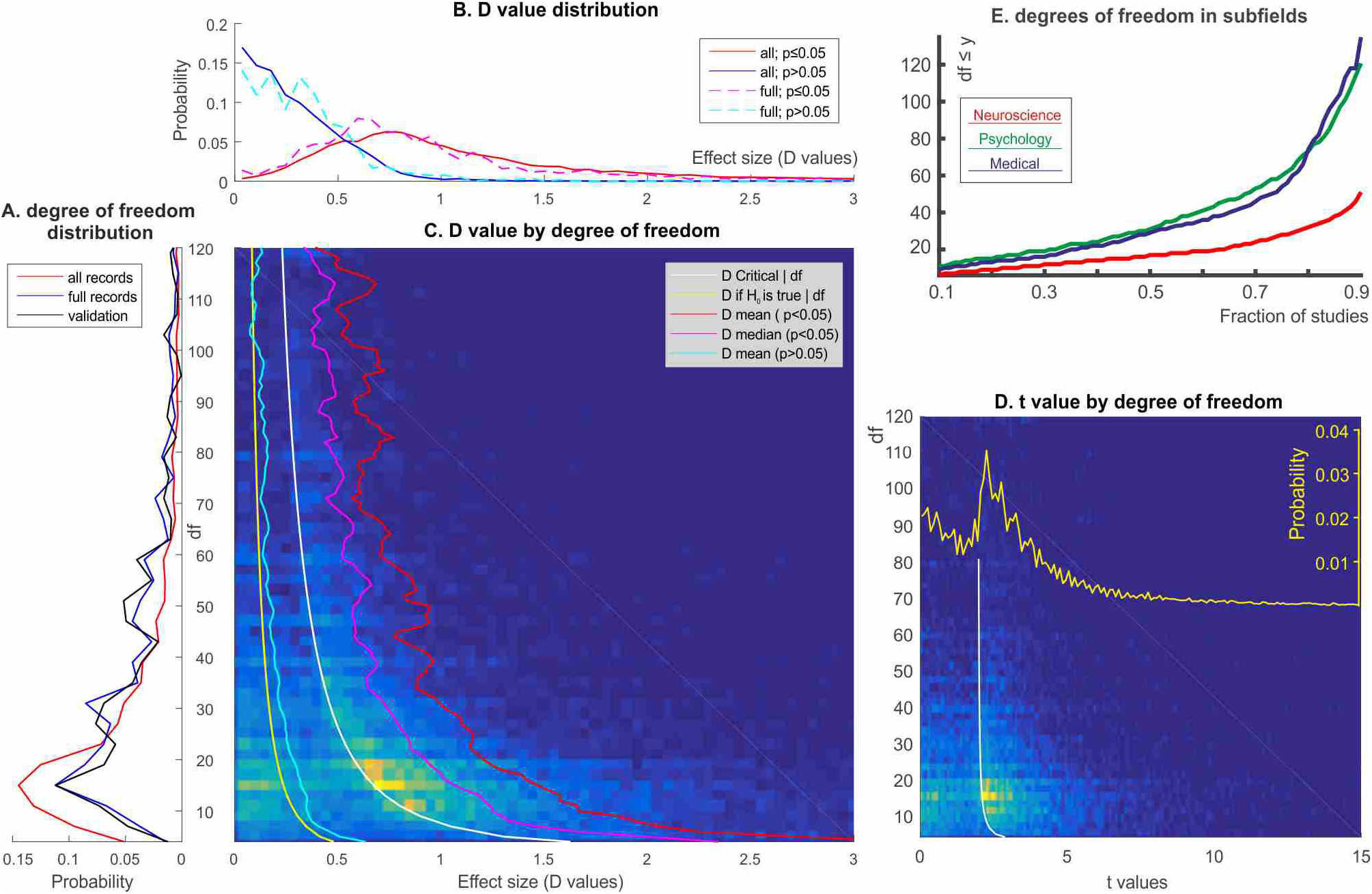
Degrees of freedom (df), effect size (D value) and t value distribution in the literature. The histograms are zoomed in for better visibility but all data were used in calculations. Colour schemes reflect the probability of observations and are scaled to the minimum and maximum values in the images. **(A)** The distribution of degrees of freedom in all records, in records with D values (full records) and in manually validated records. **(B)** The distribution of D values in significant and non-significant (p>0.05) records in the whole data set and in the subset of data with D value reports (full records). **(C)** The bivariate distribution of D values with degrees of freedom in the whole data set. The white curve denotes the significance threshold for α≤0.05 in terms of D values. The red and magenta curves show the mean and median of statistically significant effect sizes only, for various degrees of freedom. The yellow curve shows the expected value of effect sizes if the null hypothesis is true. The cyan curve shows the mean effect size from non-significant records in the data (the median was nearly the same). **(D)** The bivariate distribution of t values with degrees of freedom in the whole data set. The yellow curve overlays the univariate distribution of t values. Note how publishing only overwhelmingly significant reports distorts the observed distribution of t values. **(E)** The fraction of studies with at least a certain level of degrees of freedom by science subfield.

**Fig. 1C.** shows the bivariate distribution of D values and degrees of freedoms and represents the mean and median effect sizes for statistically significant and non-significant records and for the whole data-set. The large discrepancy between statistically significant and non-significant results is clear (medians, 25th and 75th quantiles for statistically significant and non-significant D values, respectively: d=0.932 [0.637-1.458]; d=0.237 [0.106-0.421]). **Fig. 1D.** shows the t value distribution associated with effect sizes. It is evident that a very large proportion of non-significant results are not reported at all (compare the shape of the distribution to the expected shapes shown in **Supplementary Figure 1B** and also see **Supplementary Figure 2B** for the distribution of p values).

For a certain effect size power is determined by sample size which determines degrees of freedom. The median of degrees of freedom was 20 for statistically significant and 19 for statistically non-significant results (mode=15 for both). Subfields showed large differences in degrees of freedom with most records having much lower degrees of freedom in cognitive neuroscience than in psychology and medicine (**Fig 1E.** 25th and 75th centiles for all records for cognitive neuroscience journals: df=10-28; psychology: df=17-60; medical journals: df=15-54). Reported effect sizes also differed markedly by subfields (25th and 75th centiles for all records for cognitive neuroscience journals: d=0.34-1.22; psychology: d=0.29-0.96; medical journals: d=0.23-0.91). The larger reported effect sizes in cognitive neuroscience may well be the consequence of effect size inflation due to having less degrees of freedom as the following power analyses suggest.

Median degrees of freedom and median effect sizes for each journal are depicted in **Fig 2A.** The blue crosses on the left show values for non-significant records. The red (cognitive neuroscience), green (psychology) and orange (medical) crosses show effect sizes for statistically significant records. Taking into account the reported degrees of freedom we computed power (at α=0.05) for effect sizes which are traditionally considered low (d=0.2), medium (d=0.5) and high (d=0.8) (Rossi, 1990; Sedlmeier and Gigerenzer, 1989; Cohen, 1962).

**Figure 2.**
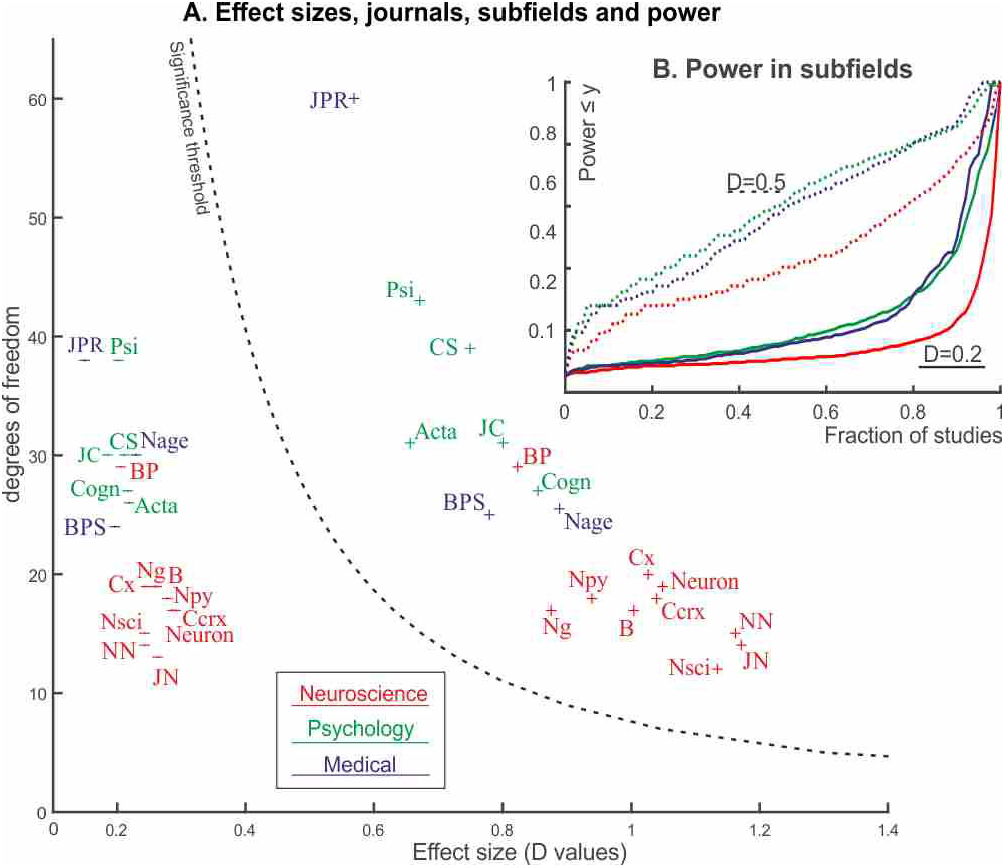
Power in journals and subfields. **(A)** Median effect sizes and degrees of freedoms in the journals analysed colour coded by science subfields. Values for non-significant data are shown on the left of the significance threshold line by horizontal lines (range of medians: D=0.15-0.29).Values for statistically significant data are shown on the right of the significance threshold line by plus signs (range of medians: 0.57-1.17). **(B)** The fraction of studies with at least a certain level of estimated power by science subfield. Journal abbreviations: Neuroscience: Nature Neuroscience (NN), Neuron, Brain (B), The Journal of Neuroscience (JN), Cerebral Cortex (Ccrx), NeuroImage (Ng), Cortex (Cx), Biological Psychology (BP), Neuropsychologia (NPy), Neuroscience (NSci). Psychology: Psychological Science (Psi), Cognitive Science (CS), Cognition (Cogn), Acta Psychologica (Acta), Journal of Experimental Child Psychology (JC). Medically oriented journals: Biological Psychiatry (BPS), Journal of Psychiatric Research (JPR), Neurobiology of Ageing (Nage).

The cumulative probability of records reaching a certain level of power is shown in **Fig. 2B.** Median and mean power are shown in **Table 1.** It is apparent that cognitive neuroscience studies had the lowest level of power. For example, to detect a small effect (d=0.2), 90% of cognitive neuroscience records had power < 0.234 which is much worse chance to detect a true effect than relying on throwing up a coin (Sedlmeier and Gigerenzer, 1989). Importantly, the somehow higher power in the journals we classified as more medically oriented was driven by the Journal of Psychiatry Research (JPR in **Fig. 2A;** median power to detect small, medium and large effects: 0.23, 0.74, 0.86) which includes more behavioural studies than the other two journals we classified as ‘medical’. These other two journals, more focused on neuroimaging, still performed better than cognitive neuroscience journals and at about the same level as psychology journals (median power to detect small, medium and large effects: 0.14, 0.53, 0.78).

**Table 1.** Median and mean power to detect small, medium and large effects in the current study and in three often cited historical power surveys. The bottom row shows mean power computed from 25 power surveys.

| Subfields or other Surveys | Records/Articles | Small effect |  | Medium effect |  | Large effect |  |
| --- | --- | --- | --- | --- | --- | --- | --- |
|  |  | Median | Mean | Median | Mean | Median | Mean |
| **Cognitive Neuroscience** | 7888/1192 | 0.11 | 0.14 | 0.40 | 0.44 | 0.70 | 0.67 |
| **Psychology** | 16887/2261 | 0.16 | 0.23 | 0.60 | 0.60 | 0.81 | 0.78 |
| **Medical** | 2066/348 | 0.15 | 0.23 | 0.59 | 0.57 | 0.80 | 0.77 |
| **All subfields** | 26841/3801 | 0.11 | 0.17 | 0.44 | 0.49 | 0.73 | 0.71 |
| **Cohen (1962)** | 2088/70 | 0.17 | 0.18 | 0.46 | 0.48 | 0.89 | 0.83 |
| **Sedlmeier & Gigerenzer (1989)** | 54 articles | 0.14 | 0.21 | 0.44 | 0.50 | 0.90 | 0.84 |
| **Rossi (1990)** | 6155/221 | 0.12 | 0.17 | 0.53 | 0.57 | 0.89 | 0.83 |
| **Rossi (1990); means of surveys** | 25 surveys |  | 0.26 |  | 0.64 |  | 0.85 |

A comparison to prominent older surveys of power estimates 25 and >50 years ago showed that median power in psychology journals have increased to detect medium sized effects but remained about the same for small and large effects (see **Table 1**; Rossi, 1990; Sedlmeier and Gigerenzer, 1989; Cohen, 1962). Power for cognitive neuroscience and for all subfields together was lower than median and mean power reported in 1962, more than half a century ago^9^. Journal impact factors negatively correlated with median power (correlation for small, medium and large effect sizes, respectively with 95% accelerated and bias corrected bootstrap confidence intervals [10^5^ permutations]: r=-0.42 [-0.63; -0.09]; -0.46 [-0.71; -0.09]; -0.45 [-0.77; -0.02]) due to the fact that cognitive neuroscience journals had the highest impact factors in our sample, but the worst power estimates.

FRP depends on power, the odds of true H_0_ to H_1_ data and on reporting bias (Ioannidis, 2005; Berger and Sellke, 1987). The continuous lines in **Fig. 3** estimate lower limits for FRP, using the probably highly inflated effect sizes computed from published t values to calculate power, for various H_0_:H_1_ odds and bias values and for α = 0.05. In the best but unrealistic case of having H_0_:H_1_ odds=1:1=1 and zero bias, FRP is 13.5%. A 10% bias pushes this to 23%. Staying in the still optimistic zone when every second to every 6th of hypotheses work out (1 ≤ H_0_:H_1_ odds ≤ 5) and with relatively modest 10-30% experimenter bias, FRP is 23-71% (median = 51%). That is, between one to three quarters of statistically significant results will be false positives. If we now move into the domain of more exploratory research where even more experimental ideas are likely to be false (5 < H_0_:H_1_ odds < 20; bias = 10-30%) then FRP grows to at least 60-91% (median = 77%).

**Figure 3.**
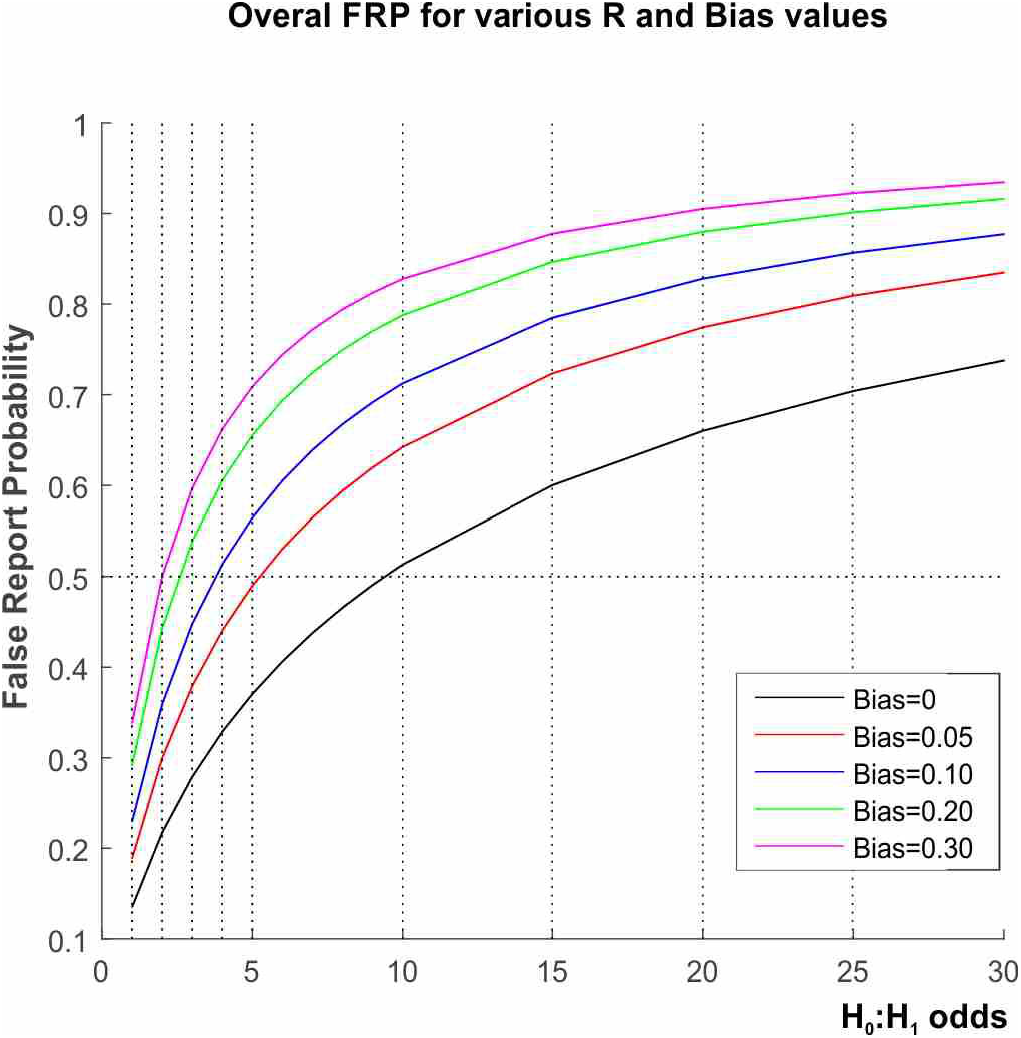
Lower estimates of false report probability (FRP) for various H_0_:H_1_ odds and Bias values.

When comparing reported to computed p values from F tests, we found that in 2.3% (n=1546 of 68,762) of all reported results with original p value reports from 14% of papers (879 papers), the reported p values were statistically significant, but when we computed the p values given the provided degrees of freedom and F statistic, these were actually nonsignificant (e.g. a study gave 2 and 45 as nominator and denominator degrees of freedom for an F-test statistic of 3.13 and reported the p-value as p≤0.05, while with these values, the p-value is p=0.0534). This finding is very similar to previous such estimates (Bakker and Wicherts, 2011; Nuijten et al. 2015; Veldkamp et al. 2014). In 10% of the above erroneous reports the p value error was larger than 0.01 and in about 20% of cases it was larger than 0.006 (see **Supplementary Figure 2)**. In 15% of papers with reporting errors affecting nominal significance more than 25% of all p value records were erroneous so that they were significant rather than non-significant. Conversely, in 0.7% (n=487) of all reported results from 323 papers, the computed p values were significant but reported p values were non-significant. All journals and all three science subfields were equally affected by reporting errors and both the proportion of articles with more favorable p value records (r=0.635 [95% bootstrap confidence interval: 0.17-0.87]) and the proportion of more favourable p value records (r=0.572 [95% bootstrap confidence interval: 0.10-0.93]) positively correlated with journal impact factors (see further details and results in **Supplementary Material, Section 5**).

## General discussion

The probability of research hypotheses depends on H_0_:H_1_ odds, experimenter bias and power (Pollard and Richardson, 1987; Berger and Sellke, 1987; Ioannidis, 2005). H_0_:H_1_ odds are inherent to each research field, and some biases are more recalcitrant and difficult to control than others, while power can in principle be easily increased by increasing sample size. However, contrary to its importance for the economic spending of research funding, the accurate estimation of effect sizes and minimizing FRP our data suggest that power in cognitive neuroscience and psychology papers is stuck at an unacceptably low level. Overall power has not improved during the past half century (Cohen 1962; Sedlmeier and Gigerenzer, 1989; Rossi, 1990). Results are also similar to other fields, such as behavioural ecology where power to detect small and medium effects was 0.13-0.16 and 0.4-0.47, respectively (Jennions and Moller, 2003).

We actually found that cognitive neuroscience journals had much lower power levels than more psychologically and medically oriented journals. This confirms previous similar inference asserting that FRP is likely to be high in the neuroimaging literature (Button et al. 2013; Yarkoni 2009). This phenomenon can appear for a number of reasons. First, neuroimaging studies and other studies using complex and highly sophisticated measurement tools in general tend to require more expensive instrumentation than behavioural studies and both data acquisition and analysis may need more time investment and resources per participant. This keeps participant numbers low. A related issue is that science funders may have reluctance to fund properly powered but expensive studies. Second, data analysis is highly technical and it can be very flexible, and many analytical choices have to be made on how exactly to analyse the results. This allows for running a very high number of undocumented and sometimes poorly understood and difficult to replicate idiosyncratic analyses influenced by a large number of arbitrary ad hoc decisions which, in their entirety, may be able to generate statistically significant false positive results with high frequency (Carp 2012; Vul et al. 2009; Kriegeskorte, 2009; Uttal 2012), especially when participant numbers are low. It is also important to consider that complicated instrumentation and (black box) analysis software is now more available but training may not have caught up with this wider availability. Third, in relation to more medical journals the stakes at risk are probably lower in cognitive neuroscience (no patients will die, at least not immediately) which may also allow for more biased publications. Nevertheless, generalizations need to be cautious, since there can be large variability in the extent of these potential biases within a given subfield. Some teams and subfields may have superb, error-proof research practices, while others may have more frequent problems. The power failure of the cognitive neuroscience literature is even more notable as neuroimaging (‘brain based’) data is often perceived as ‘hard’ evidence lending special authority to claims even when they are clearly spurious (Weisberg, 2008).

The reporting of p values as significant when they are not (even when juxtaposed against the other statistics form the same analysis) are probably only a modest contributor in the greater drive to generate and report statistically significant results. Most of the significance chasing would not be possible to detect through such inconsistencies. The high prevalence of reported significant results (amounting to three quarters of the reported p values and related statistics) it unrealistically high and consistent with strong selective reporting biases. This has been demonstrated also in distributions of p values reported in abstracts and full texts of biomedical papers (Chavalarias, et al. 2016). The positive correlation between p value reporting errors and journal impact factors suggests that such errors are more prevalent in ‘high impact’ journals and these journals also had less power in the studies that they reported.

Some limitations need to be mentioned for our study. First, given the large scale automation, we cannot fully verify whether the extracted data reflect primary, secondary or even trivial analyses in each paper. In the absence of pre-registered protocols, however, this is extremely difficult to judge even when full papers are examined. Evaluation of biomedical papers suggests that many reported p-values, even in the abstracts, are not pertinent to primary outcomes (Ioannidis 2014). Second, some types of errors, such as non-differential misclassification, may lead to deflated effect sizes. However, in the big picture, with very small power, inflation of the statistically significant effects, is likely to be more prominent than errors reducing the magnitude of the effect size. Third, given the large scale automated extraction, we did not record information about characteristics of the published studies, e.g. study design. It is likely that studies of different designs (e.g. experimental versus observational studies) may have different distribution of effect sizes, degrees of freedom, and power even within the same sub-discipline.

In all, the combination of low power, selective reporting and other biases and errors that we have documented in this large sample of papers in cognitive neuroscience and psychology suggest that high FRP are to be expected in these fields. The low reproducibility rate seen for psychology experimental studies in the recent Open Science Collaboration (Nosek et al. 2015a) is congruent with the picture that emerges from our data. Our data also suggest that cognitive neuroscience may have even higher FRP rates, and this hypothesis is worth evaluating with focused reproducibility checks of published studies. Regardless, efforts to increase sample size, and reduce publication and other biases and errors are likely to be beneficial for the credibility of this important literature.

## Materials and Methods

### Data extraction and validation

We extracted statistical information from cognitive neuroscience and psychology papers published as pdf files. We sampled a range of 18 journals frequently cited in cognitive neuroscience and psychology. Our aim was to collect data on the latest publication practices. To this end, we analysed four years’ regular issues of all journals published between Jan 2011 to Aug 2014.

We categorized 10 journals as focused more on (cognitive) neuroscience (Nature Neuroscience, Neuron, Brain, The Journal of Neuroscience, Cerebral Cortex, NeuroImage, Cortex, Biological Psychology, Neuropsychologia, Neuroscience) and 5 journals focused more on psychology (Psychological Science, Cognitive Science, Cognition, Acta Psychologica, Journal of Experimental Child Psychology). We also searched three more medically oriented journals which are nevertheless often cited in cognitive neuroscience papers so as to increase the representativeness of our sample (Biological Psychiatry, Journal of Psychiatric Research, Neurobiology of Ageing). Journal impact factors ranged from 2.367 (Acta Psychologica) to 17.15 (Nature Neuroscience). 5-year impact factors were considered as reported in 2014 (see **Supplementary Table 1**).

We extracted statistical information about t tests and F tests (t values, F values, degrees of freedoms, p values and effect sizes). A computer algorithm searched through each paper for frequently occurring word and symbol combinations for reporting degrees of freedoms and effect sizes provided as Cohen’s D. When there were less than 20 empirical papers in a journal issue all empirical research reports with any t and F statistics reports were analysed. When there were more than 20 papers in an issue, a random sample of 20 papers were analysed merely because this was the upper limit of papers accessible in one query. This procedure sampled most papers in most issues and journals. All algorithms and computations were coded in Matlab © 2015b.

The whole database was checked for errors in reporting and potential double entries so that only unique records were kept for analysis. Further, during a validation procedure we randomly selected 100 papers with t value, df and effect size reports and 100 papers with F value and df reports. We manually checked through all records in these papers and compared them to the automatically compiled database. This was done to see the accuracy of the computer algorithm and to gather information on the data. Validation results showed that the automatic extraction mechanism performed very well. The papers selected for validation included 1478 records of data, the algorithm missed only 76 records (5.14%), usually due to atypical punctuation or line breaks. We found that the overwhelming majority of records related to two sample t-tests reported close to equal group numbers (median ratio of group numbers = 1). The ratio of the participant numbers in the larger group to the participant numbers in the smaller group was smaller than 1.15 in 77% of records. We also established that with a df value of 10 or less about 94% of tests were one sample or matched t-tests whereas about 72% of records with higher df values were one-sample t-tests. The F test validation procedure for 100 randomly selected papers showed that the algorithm missed only 84 out of 1084 records (92.25% of records correctly found).

### Computing effect sizes from t tests

t-test data was used for effect size, power and false report probability analysis as it is straightforward to estimate effect sizes from published t values. After checks for reporting errors 7 records with df>10,000 were excluded from analysis as clear outliers. This left 27,414 potential records. 26,841 of these records from 3801 papers had both degrees of freedoms and t values reported. We used this data for the effect size analysis. 17,207 t-test records (64.1%) were statistically significant (p≤0.05) and 9634 (35.9%) t-test records were statistically nonsignificant (p>0.05). 2185 t-test records also reported Cohen's D as a measure of effect size (1645 records with p≤0.05 [75.3%] and 540 records with p>0.05 [24.7%]).

As it is not possible to establish the exact participant numbers in groups for our large sample size making a few reasonable assumptions is inevitable. First, based on our validation data from 1478 records we made the assumption that participant numbers in two-sample t-test groups were equal on the average. The number of participants in groups was approximated as the upwards rounded value of half the potential total number of participants in the study, i.e.: N_subgroup_ = round_upper_((df+2)/2). This formula even slightly exaggerates participant numbers in groups, so it can be considered generous when computing power. Second, regarding matched t-tests we assumed that the correlation between repeated measures was 0.5. In such a case the effect sizes can be approximated in the same way for both one-sample and matched t-tests. These assumptions allowed us to approximate effect sizes associated with all t-tests records in a straightforward way (Hunter and Schmidt, 1990; Fritz et al. 2002). Computational details are provided in **Supplementary Material, Section 2.**

Considering the validation outcomes we assumed that each record with a df of 10 or less had 93% chance to be related to a one-sample or matched-sample t-test and other records had 72% chance to be related to a one-sample or matched-sample t-test. Hence, we estimated the effect sizes for each data record with an equation assuming a mixture of t-tests where the probability of mixture depended on the df value:

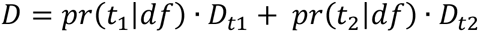

where pr(t1|df) and pr(t2|df) refer to the respective probabilities of one sample and matched t-tests (t_1_) and independent sample t-tests (t_2_) and D_t1_ and D_t2_ refer to the respective effect sizes estimated for these tests.

The power of t-tests was computed from the non-central t distribution assuming the above mixture of one-sample, matched and independent sample t-tests (Harrison and Brady, 2004). Computational details are provided in **Supplementary Material, Section 3**.

Published effect sizes are likely to be highly exaggerated. Using these exaggerated effect sizes for power calculations will then overestimate power. Hence, if we calculate FRP based on power calculated from published effect size reports we are likely to estimate the lower limits of FRP. So, we estimated the lower limits for FRP, using the probably highly inflated effect sizes (computed from published t values) to calculate power for various H_0_:H_1_ odds and bias values and with α = 0.05. (The computation of FRP is laid out in detail in **Supplementary Material, Section 4**.)

In order to get the expected value of FRP for the whole literature we weighed the FRP computed for each df and D value combination by the probability of that particular (df,D) combination occurring in the research literature and summed the results for all (df,D) combinations:

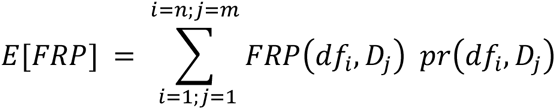

### Checking for p value reporting errors: F tests

F tests were used for examining p value reporting errors as this is more difficult with t-tests which may report either one and two-tailed p values. We extracted 75,904 records of data. 73,532 F-test records from 6273 papers were used for examining p value reporting errors. 51,920 records (70.6%) were statistically significant and 21,612 records (29.4%) were not significant. 68,762 records also explicitly reported p values (significant: 51,338 [74.7%]; nonsignificant: 17,424 [25.3%). This data allowed us to compare exact computed (from degrees of freedoms and F values) and reported p values.

## Acknowledgments

DS is supported by the James S McDonnell Foundation.

## Supplementary Material

### Section 1:Journal information

**Supplementary Table 1.** shows information about the journals examined.

**Supplementary Table 1.** Journal information for the three subfields considered for t tests. 5-year journal impact factors used in the study; the number of records in journals; the number of papers by journals and the average number of records per paper.

| **Journals** | **5 year I.F.** | **Records** | **Papers** | **Records/Papers** |
| --- | --- | --- | --- | --- |
| **A. Psychology** |  |  |  |  |
| Psychological Science | 6.16 | 2208 | 387 | 5.7 |
| Cognitive Psychology | 5.508 | 493 | 53 | 9.3 |
| Cognition | 5.088 | 2505 | 316 | 7.9 |
| Acta Psychologica | 3.016 | 769 | 161 | 4.8 |
| JECP | 3.353 | 1913 | 275 | 7.0 |
| **B. Neuroscience** |  |  |  |  |
| Nature Neuroscience | 17.15 | 1212 | 121 | 10.0 |
| Neuron | 16.485 | 834 | 79 | 10.6 |
| Brain | 10.846 | 584 | 75 | 7.8 |
| The Journal of Neuroscience | 7.87 | 5408 | 621 | 8.7 |
| Cerebral Cortex | 8.372 | 1744 | 252 | 6.9 |
| Neuroimage | 6.956 | 971 | 193 | 5.0 |
| Cortex | 5.389 | 1508 | 198 | 7.6 |
| Biological Psychology | 4.173 | 1338 | 205 | 6.5 |
| Neuropsychologia | 4.495 | 2089 | 354 | 5.9 |
| Neuroscience | 3.458 | 1199 | 163 | 7.4 |
| **C. Medical** |  |  |  |  |
| Biological Psychiatry | 10.347 | 1101 | 187 | 5.9 |
| Journal of Psychiatric Research | 4.46 | 468 | 94 | 5.0 |
| Neurobiology of Aging | 5.127 | 497 | 67 | 7.4 |

### Section 2:Computing effect sizes from t-tests

Effect size calculation for one-sample t-test:

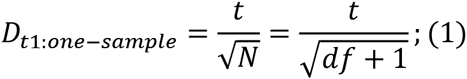

where *N* stands for the potential total number of participants which is df+1 for one-sample and matched t-tests.

Because, by definition:

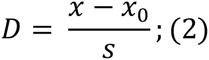

So:

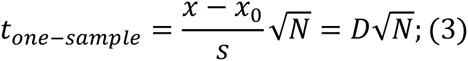

Effect size calculation for two sample t-test:

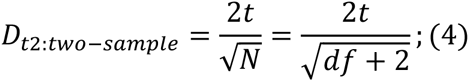

where *N* stands for the potential total number of participants which is df+2 for independent sample t-tests.

Because

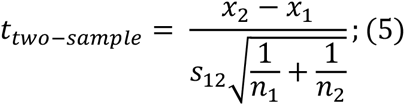

Where:

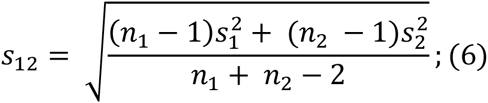

And by definition:

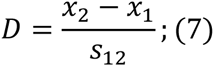

Assuming that n_1_ = n_2_ = n_c_ and letting N = n_1_+n_2_ (ie. N = total number of participants) we get:

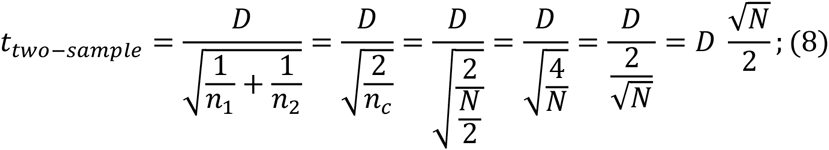

So:

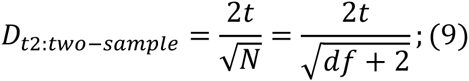

For matched-sample t-tests:

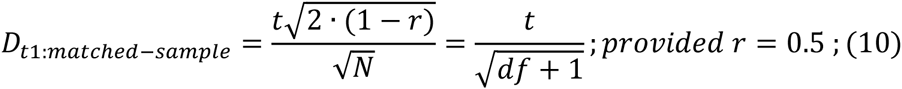

Based on our mixture model of one-sample and two-sample t-tests we can expect a t value distribution as depicted in **Supplementary Figure 1A-B.** The figure illustrates the mechanism of effect size exaggeration and the importance of H_0_:H_1_ odds.

**Supplementary Figure 1.**
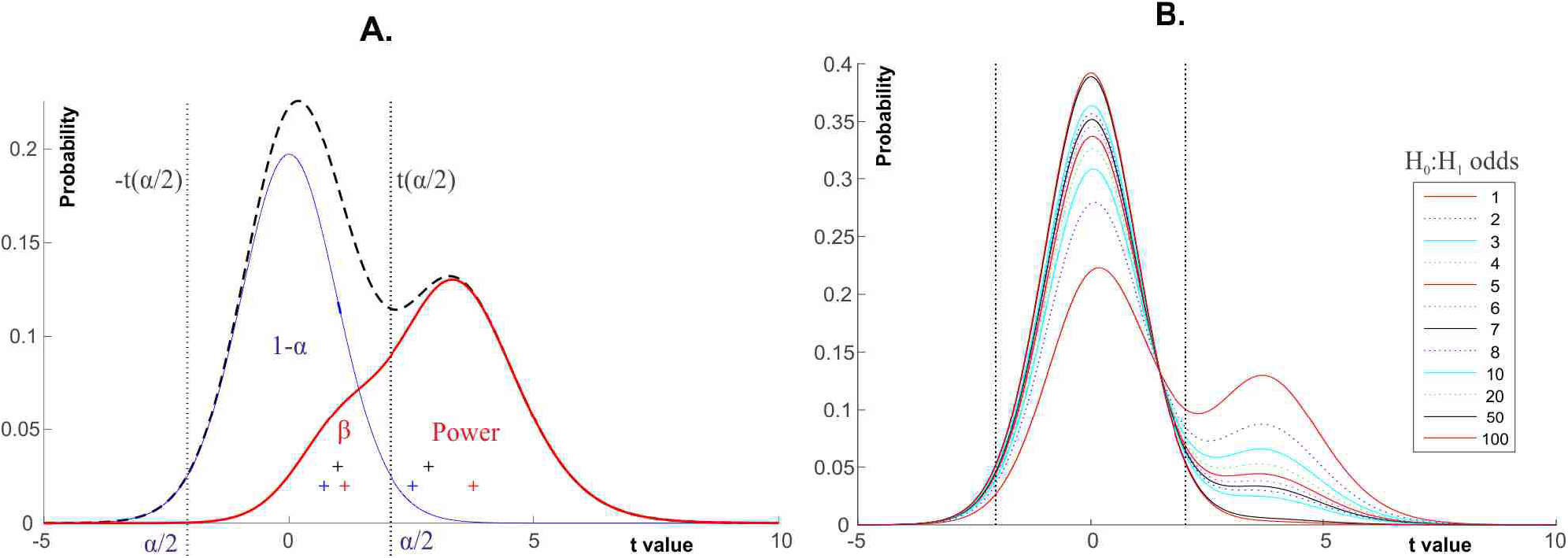
t value distributions when all negative and positive results are published. **(A)** Illustration of effect size exaggeration due to lack of power (df=22). ±t(α) stand for the critical t values with α=0.05. The figure depicts the probability density of t values under a mixture model (Eq. 11) assuming a 70% proportion of one-sample t-tests. The blue line denotes the probability density of t values if the null hypothesis is true. The red line denotes the probability density of t values if the alternative hypothesis is true with an effect size of D=0.75. The black dashed line denotes the probability density of t values if in half the data the null hypothesis is true and in the other half the alternative hypothesis is true (ie. The H_0_:H_1_ odds are 1). The crosses mark the expected value of absolute t values (which depend on the actual effect size measured), note that these are dramatically different in significant and nonsignificant data irrespective of whether the null hypothesis is true or not. Blue crosses: the expected t value in data where the null hypothesis is true and the test outcome is non-significant (left cross: true negative) and when the test outcome is significant (right cross: false positive). Red crosses: the expected t value in data where the alternative hypothesis is true and the test outcome is non-significant (left cross: false negative) and when the test outcome is significant (right cross: true positive). Black crosses: the expected t values in non-significant (left cross) and significant (right cross) data. **(B)** Expected mixture model t value distribution for df = 30 for various H_0_:H_1_ odds.

### Section 3:Power calculations

The power of t-tests can be computed from the non-central t distribution assuming an mixture of t tests (see main text for the mixture model we used).

For simpler writing we can first define the cdf for the mixture distribution where the probability of a t value depends on the proportion of one and two-sample t-tests in the data:

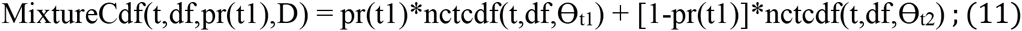

Where

nctcdf stands for the cumulative non-central t probability density function,

t stands for a t value,

df stands for the given degrees of freedom,

pr(t1) stands for the probability of a one-sample or matched t-test.

⊖_t1_ and ⊖_t2_ are non-centrality parameters dependent on Cohen's D (Harrison and Brady, 2004):

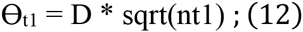

Where nt1 = total number of participants; i.e. nt1 = df+1.

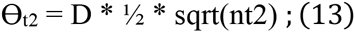

Where
nt2 = the number of participants in one experimental group; i.e. nt2 = roundupper((df+2)/2).

Given the above notation, power for a given effect size (D), df and critical t value (t(α)) can be computed as (Harrison and Brady, 2004):

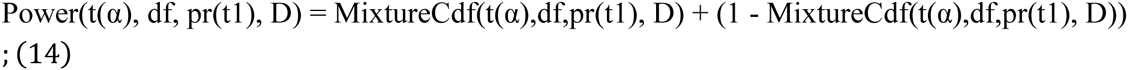

### Section 4: False and True Report probability

If we use Null Hypothesis Significance Testing (NHST) then the long run False Report Probability (FRP) is the long run probability that the null hypothesis (H_0_) is true when we get a statistically significant finding. The long run True Report Probability (TRP) is the long run probability that the alternative hypothesis (H_1_) is true when we get a statistically significant finding. Computationally, FRP is the number of statistically significant false positive findings divided by the total number of statistically significant findings. TRP is the number of statistically significant truly positive findings divided by the total number of statistically significant findings.

FRP and TRP can be computed by Bayes’ theorem. If we take ‘sig’ to stand for ‘statistically significant test outcome’ then the total probability of finding a statistically significant result is:

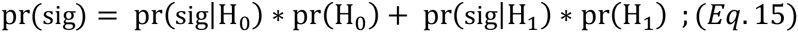

Hence, FRP and TRP can be written as:

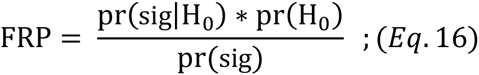

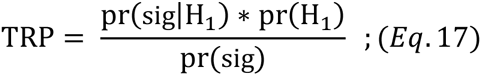

Considering a long run of NHST studies, the long run probability of having a significant test outcome when H_0_ is true is α and the long run probability of having a significant test outcome when H_1_ is true is Power = 1 - β. That is, α = pr(sig|H_0_) and Power = p(sig|H_1_). Hence, Eq.16. and Eq. 17. can be re-written as:

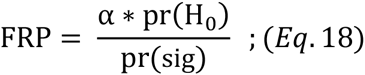

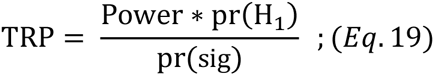

Eq.16. and Eq.17. can also be expressed in terms of odds ratios of true H_0_ and true H_1_ situations. For example, we can denote the odds of true H_0_ situations as ‘O’ and write:

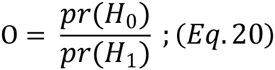

We can express pr(H_0_) using the above as:

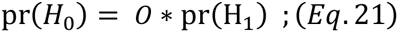

then Eq.16. Can be rewritten as:

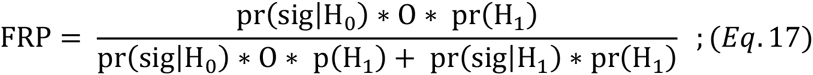

This can be simplified by pr(H_1_):

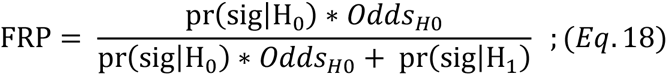

Using α and Power = 1 - β we can write:

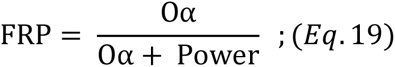

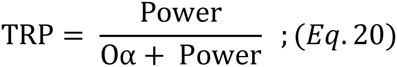

FRP + TRP = 1; e.g.:

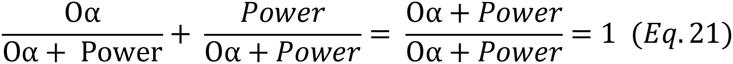

Consequently:

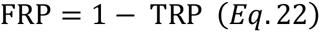

and

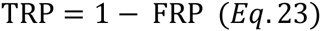

Equivalently to the above, we can also express the odds of true H_1_ situations as the ratio of pr(H_1_) and pr(H_0_) and denote it with ‘R’ as in Ioannidis (2005):

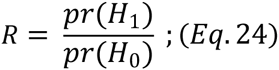

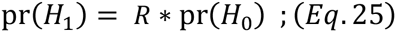

Substituting Eq.25. into Eq.16.:

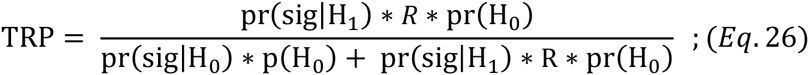

Simplifying:

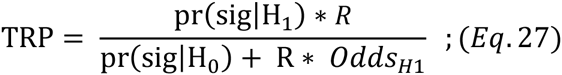

Using α and Power=1 - β we can write:

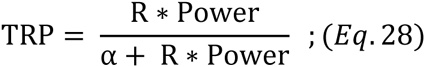

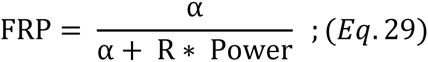

Eq.28. is equivalent to the one used by Ioannidis (2005) with a slightly different notation.

He defined PPV = TRP; Power = 1-β and equivalently to Eq. 28 he wrote:

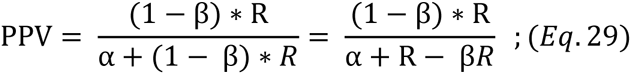

Ioannidis (2005) also defined Bias, signified by ‘*u*’. On the one hand, bias results in categorizing fraction *u* β(u*(1 - α)) of otherwise true negative results (in case there is no bias) as positive results. On the other hand, bias results in categorizing fraction *u* (uβ=u*(1 - Power)) of otherwise (in case there is no bias) missed true positive results as positive results. That is, bias alters Eq.28. as:

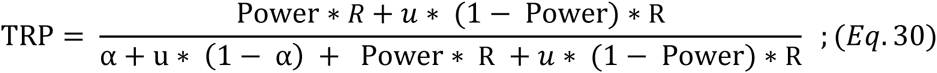

Using the notation of Ioannidis (2005) this can be rewritten as:

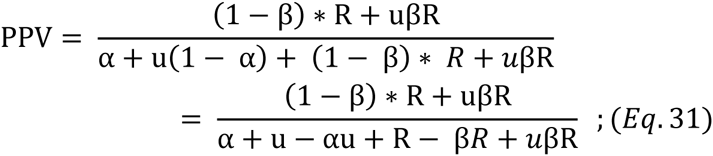

Also, notice the relation between O and R:

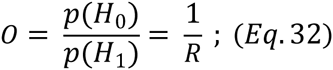

Hence,

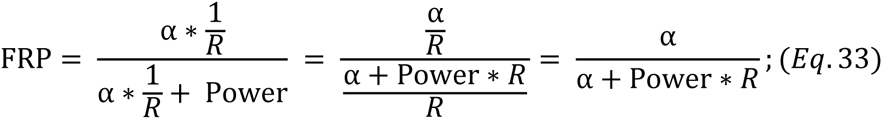

Similarly:

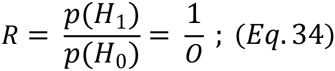

Hence,

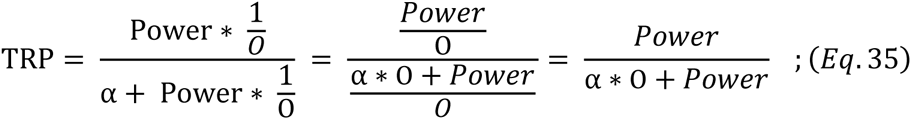

### Section 5:Supplementary Results; p value reporting errors

A very large number of records relied on graded p value reporting as shown in **Supplementary Figure 2A.** (e.g. reporting p<0.05 instead of p=0.047). There was a definite bias towards reporting significant p values (**Supplementary Figure 2B.**).

**Supplementary Figure 2C** shows the distribution of the difference between reported and computed p values. First, if computed p values were significant, relatively few records reported lower p values than the computed ones and most reported p values were in fact larger than the computed ones due to graded reporting (green curve in **Supplementary Figure 2B**). In contrast, when computed p values were non-significant there was more variability between reported and computed p values (black dashed curve in **Supplementary Figure 2C**) into both negative and positive directions, mostly in the p±0.01 range. The difference between the respective distributions is depicted by the black curve in **Supplementary Figure 2D**.

In 1546 cases (3% of all reported significant results) from 879 papers (14% of all papers) computed p values were non-significant but researchers reported significant p values (continuous red curve in **Supplementary Figure 2C**). In 10% of such cases the p value error was larger than 0.01 and in about 20% it was larger than 0.006 (see circled points in **Supplementary Figure 2C**). In such cases the absolute magnitude of the deviation from the real p value was much larger than in the case when both computed and reported p values were non-significant (blue curve in **Supplementary Figure 2C**). The difference between the respective distributions is depicted by the red curve in **Supplementary Figure 2D**.

In 2.79% of all reported non-significant results (487 cases) from 323 papers computed p values were significant but reported p values were non-significant (dotted red curve in **Supplementary Figure 2C**). Again, there was a higher proportion of p values which were highly diminished relative to their real value compared to the distribution of p value differences in the case when both the computed and reported p values were non-significant (dotted blue curve in **Supplementary Figure 2C** and see the dotted red difference curve in **Supplementary Figure 2D**).

**Supplementary Figure 2.**
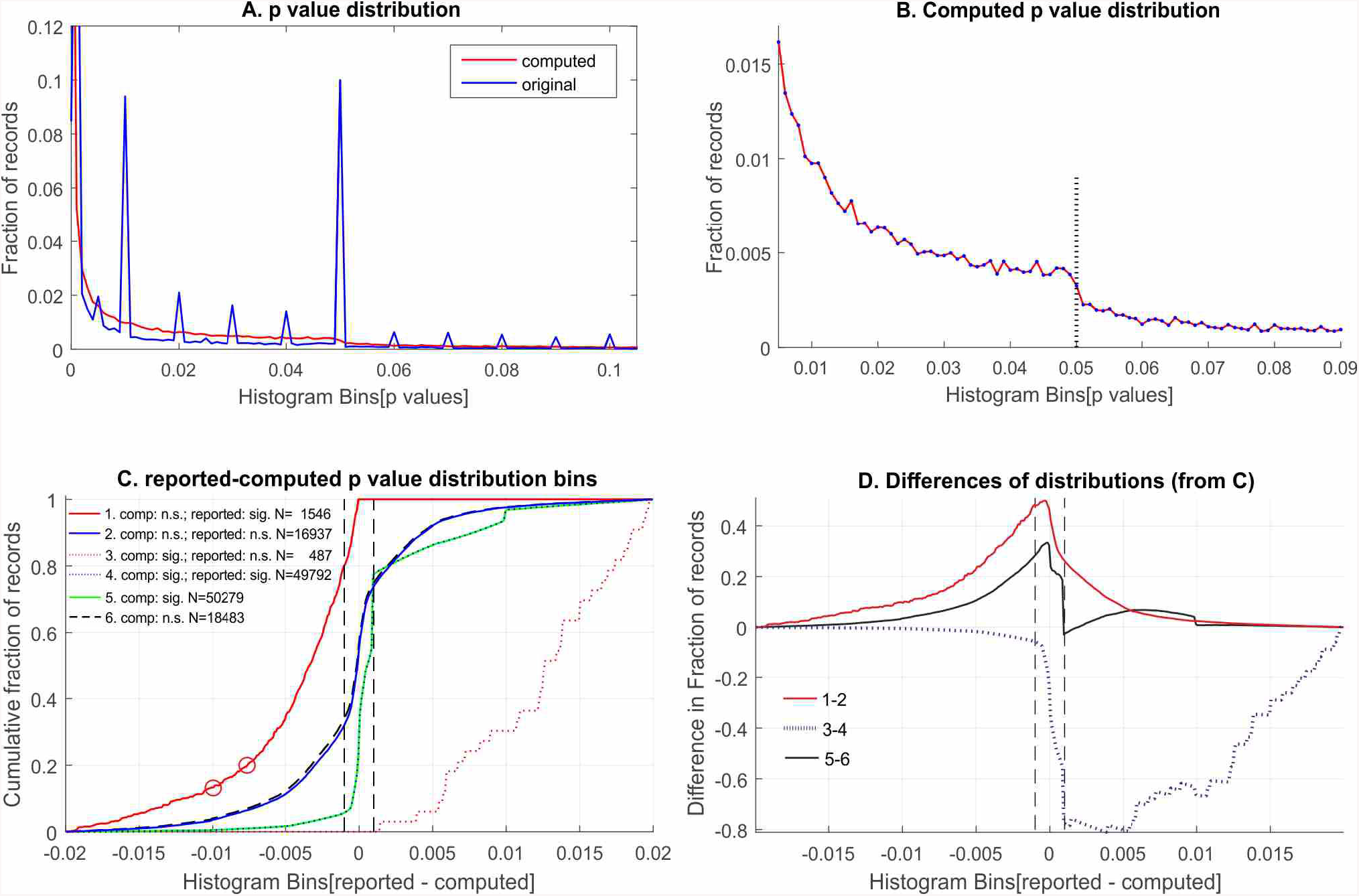
P value analysis from F tests. **(A)** Graded reporting of p values. **(B)** The distribution of computed p values reflects a clear bias to report more significant than non-significant results. **(C)** The difference between reported and computed p values (reported minus computed). See explanation in text. **(D)** The difference between the noted numbered difference distributions in **Panel B.** See explanation in text. The dashed vertical lines in Panels C and D denote potential tolerance bands for computational errors. Within such bands the differences quickly drop to zero.

**Supplementary Figure 3.**
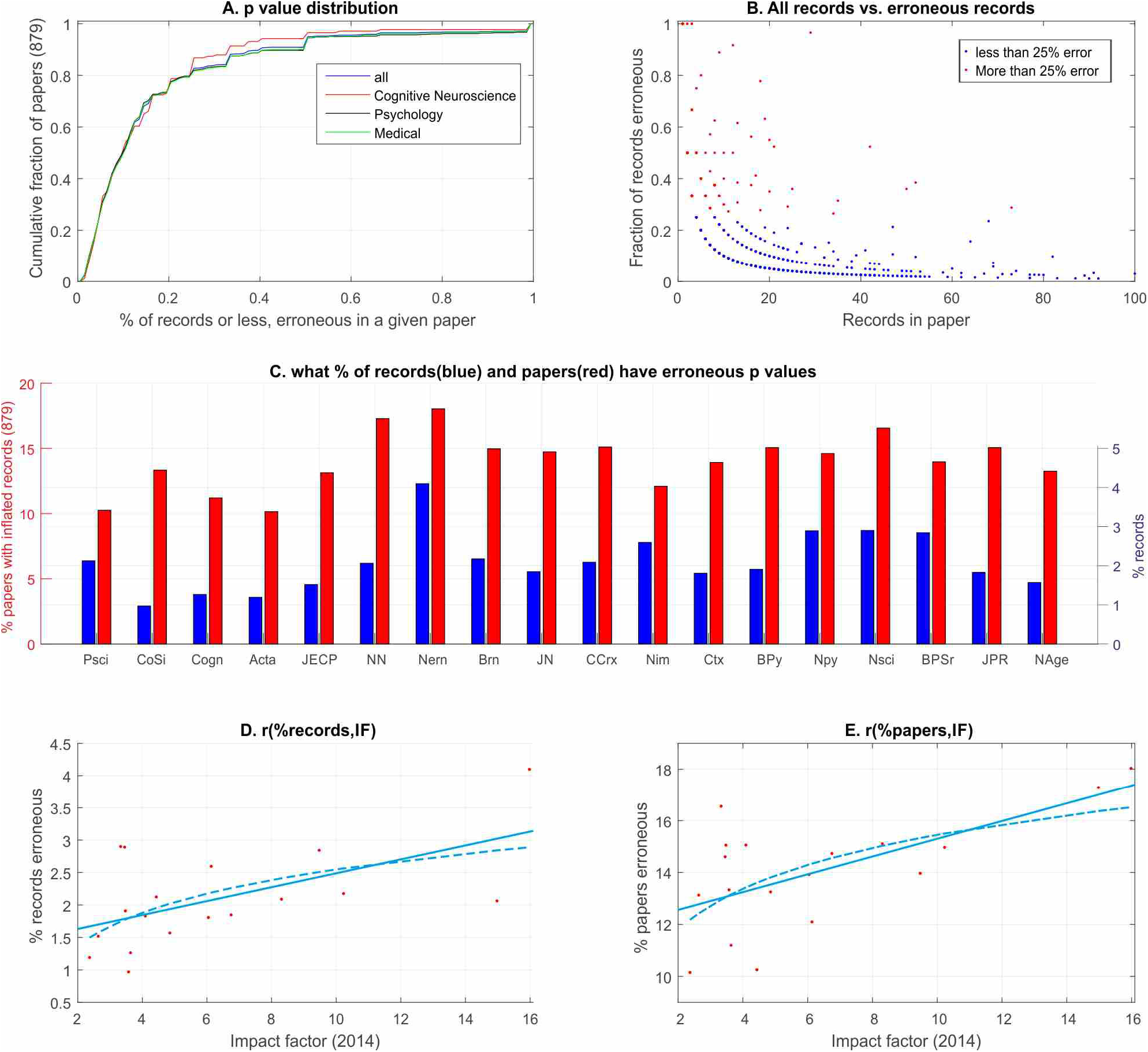
The percent of erroneous records in journals. **(A)** The fraction of the 879 papers with erroneous effect sizes (y) in which at least a certain fraction of records were erroneous (x). **(B)** The fraction of erroneous records in a paper vs. all the records in a paper. Papers with more than 25% erroneous p value records are marked by red. **(C)** The median percentage of records (blue) and papers (red) with erroneous effect sizes in a journal. **(D)** The correlation of the median percentage of erroneous record.s in papers in a given journal and journal impact factors. **(E)** The correlation of the median percentage of papers with erroneous records in a given journal and journal impact factors. Both linear and logarithmic (log[journal impact factor]) trend lines are shown.

We also assessed the fraction of records with erroneous (‘erroneous’ referring to misclassified statistical significance here) p values in all the 879 papers with erroneous p values (**Supplementary Figure 3A**). Profiles did not differ between science subfields which suggests that p value reporting errors may be equally typical of many fields. Overall, in about 50% of papers less than 10% of all reported records were erroneous but in some papers a very large number of records were erroneous (**Supplementary Figure 3B**). While a few reporting errors could be attributed to random errors it is notable that in 15% of the 879 papers with errors more than 25% of all p value records were erroneous.

We found that all journals in our sample were affected by p value inflation both by looking at the fraction of records with erroneous p values and the fraction of papers with erroneous p values (**Supplementary Figure 3C.**). Both the proportion of articles with erroneous p value records (r=0.635 [95% bootstrap confidence interval: 0.17-0.87]) and the proportion of erroneous p value records (r=0.572 [95% bootstrap confidence interval: 0.10-0.93]) positively correlated with journal impact factors (**Supplementary Figure 3D-E**).

## References

Bakker M, Wicherts JM. The (mis)reporting of statistical results in psychology journals. Behav Res Methods. 2001. 43, 666–78.

Berger JO, Sellke T. Testing a Point Null Hypothesis: the Irreconcilability of *p*-Values and Evidence. Journal of the American Statistical Association. 2001. 82, 112–122.

Button KS, Ioannidis J, Mokrysz C, Nosek BA, Power failure: Why small sample size undermines the reliability of neuroscience. Nature Reviews Neuroscience. 2013. 14, 365–376.

Carp J. The secret lives of experiments: methods reporting in the fMRI literature. NeuroImage. 2012. 63, 289–300. http://dx.doi.org/10.1016/j.neuroimage.2012.07.004

Chavalarias D, Wallach J, Li A, Ioannidis JP Evolution of reporting P-values in the biomedical literature, 2016, 1990–2015. JAMA.

Cohen J. The statistical power of abnormal - social psychological research: A review. Journal of Abnormal and Social Psychology. 1962. 65, 145–153.

Fritz CO, Morris, PE, Richler JJ. Effect size estimates: Current use, calculations and interpretation. Journal of Experimental Psychology: General. 2012. 141, 2–18.

Gigerenzer G, Marewski JN (2015), Surrogate science: The idol of a universal method for scientific inference. Journal of Management, 2015. 41, 421–440

Gigerenzer G, Swijtnik Z, Porter T, Daston L, Beatty J, Kruger L (1989), The empire of chance: How probability changed science and everyday life. Cambridge University Press. 1989

Hallahan M. Rosenthal R. Statistical Power: Concepts, procedures and applications. Behavioural Research Therapy. 1996. 34, 489–499.

Harrison DA, Brady AR, Sample size and power calculations using the noncentral tdistribution. 2004. The Stata journal. 4, 142–153.

Hunter JE, Schmidt FL. Methods of meta-analysis: Correcting error and bias in research findings. Newbury park. Sage. 1990.

Ioannidis JPA. Why most published research findings are false. PLoS Medicine. 2005. 2, e124.

Ioannidis, JPA. (2008). Why most discovered true associations are inflated. Epidemiology. 2008. 19, 640–648.

Ioannidis JPA. (2014). How to make more published research true. PLoS Medicine. 2014. 11(10): e1001747.

Ioannidis JPA. Greenland S, Hlatky MA, Khoury MJ, Macleod MR, Moher D, Schulz KF, Tibshirani R. Increasing value and reducing waste and research design, conduct and analysis. Lancet, 2014. 383, 166–175.

Jennions MD, Moller AP. A survey of the statistical power of research in behaicoural ecology and animal behaviour. Behavioural Ecology. 2003. 14, 438–445

Kriegeskorte N, Simmons WK, Bellgowan PSF, Baker CI. Circular analysis in systems neuroscience – the dangers of double dipping. Nature Neuroscience. 2009. 12, 535–40.

Nosek BA et al. (2015a). Estimating the reproducibility of psychological science. Science. 2015a. 349, 943.

Nosek BA, Spies JR, Motyl M. Scientific utopia II: Restructuring incentives and practices to promote truth over publishability. Perspectives on Psychological Science. 2013. 7, 615–631.

Nosek BA, Alter G, Banks GC, Borsboom D, Bowman SD, Breckler SJ, Buck S, Chambers CD, Chin G, Christensen G, Contestabile M, Dafoe A, Eich E, Freese J, Glennerster R, Goroff D, Green DP, Hesse B, Humphreys M, Ishiyama J, Karlan D, Kraut A, Lupia, A, Mabry, P, Madon, TA, Malhotra, N, Mayo-Wilson, E, McNutt, M, Miguel, E, Paluck, EL, Simonsohn, U, Soderberg, C, Spellman, BA, Turitto, J, VandenBos, G, Vazire, S, Wagenmakers, EJ, Wilson, R, Yarkoni, T. Promoting an open research culture. Science. Jun. 2015b. 26;348(6242):1422–5

Nuijten MB, Hartgerink CH, van Assen MA, Epskamp S, Wicherts, JM. The prevalence of statistical reporting errors in psychology (1985–2013). Behav Res Methods. 2015.

Pollard P, Richardson JTE. On the probability of making Type-I errors. Psychological Bulletin, 1987. 102, 159–163.

Rossi JS. Statistical power of psychological research: What have we gained in 20 years? Journal of consulting and clinical psychology. 1990. 58, 646–656. http://dx.doi.org/

Schmidt FL. What do data really mean? Research findings, meta-analysis and cumulative knowledge in psychology. American Psychologist, 1992. 47, 1173–1181.

Sedlmeier P, Gigerenzer G. Do studies of statistical power have an effect on the power of the studies? Psychological Bulletin. 1989. 105, 309–316.

Uttal WR. Reliability in cognitive neuroscience: A meta-meta-analysis. MIT Press.

Veldkamp CL, Nuijten MB, Dominguez-Alvarez L, van Assen MA, Wicherts JM. Statistical reporting errors and collaboration on statistical analyses in psychological science. PLoS One. 2014. 9, 1–19.

Vul E, Harris C, Winkielman P, Pashler H. Puzzlingly high correlations in fMRI studies of emotion, personality and social cognition. Perspectives on Psychological Science. 2009. 4, 274–324.

Weisberg DS, Keil FC, Goodstein J, Rawson E, Gray JR. The seductive allure of neuroscience explanations. The Journal of Cognitive Neuroscience, 2008. 20, 470–477

Yarkoni T. Big correlations in little studies: Inflated fMRI correlations reflect low statistical power—Commentary on Vul et al. Perspectives on Psychological Science. 2009. 4, 294–298

## References

Harrison D.A., Brady A.R., Sample size and power calculations using the noncentral tdistribution. The Stata journal. 4, 142–153 (2004).

Fritz, C.O., Morris, P.E., Richler, J.J, Effect size estimates: Current use, calculations and interpretation. Journal of Experimental Psychology: General. 141, 2–18 (2012).

Ioannidis, J.P.A. Why most published research findings are false. PLoS Medicine. 2, e124. (2005).

Hunter, J.E & Schmidt F.L. Methods of meta-analysis: Correcting error and bias in research findings. Newbury park. Sage. (1990).

